# Cannabidiol attenuates chemotherapy-induced peripheral neuropathic pain through a mechanism that requires the enzyme *N*-acylphosphatidylethanolamine-specific phospholipase D (NAPE-PLD)

**DOI:** 10.64898/2026.06.08.730909

**Authors:** Carlos Henrique Alves Jesus, Ailing Li, Serge Luquet, Ken Mackie, Andrea G. Hohmann

## Abstract

Cannabidiol (CBD) is a non-psychoactive component of cannabis that has been studied as a potential therapy for chronic pain. CBD attenuates behavioral hypersensitivities in models of neuropathic pain, and promotes production of bioactive lipids (e.g., anandamide), altering lipid signaling. However, a lack of understanding of the mechanisms underlying the therapeutic effects of CBD has hindered development and application of CBD to mechanism-based therapies for pain in people. We asked whether the analgesics effects of CBD were dependent upon the enzyme NAPE-PLD. We used a mouse model of chemotherapy-induced peripheral neuropathy (CIPN) to evaluate the acute and chronic antinociceptive effects of CBD and investigate its mechanisms. Pharmacological specificity was tested with antagonists targeting CB1, CB2, PPARγ, and PPARα receptors. Mechanisms were further examined using NAPE-PLD and GPR55 knockout mice. We also assessed repeated CBD dosing during both the development and maintenance of paclitaxel-induced CIPN in wild-type, GPR55 KO, and NAPE-PLD KO mice. CBD suppressed paclitaxel-induced behavioral hypersensitivities; these effects were attenuated by a PPARα and PPARγ antagonists, but not CB1 or CB2 antagonists. CBD reduced both the development and maintenance of neuropathic nociception in a model CIPN in wild-type mice, but these effects were absent in NAPE-PLD KO mice. By contrast, anti-allodynic efficacy of CBD was fully preserved in GPR55 KO mice. Pharmacological blockade of the PPARα receptor and genetic deletion of NAPE-PLD abolished the antinociceptive effects of CBD in a model of CIPN, suggesting a pivotal role for NAPE-PLD and PPAR receptors in CBD-mediated analgesia in chemotherapy-induced neuropathic pain.

## 1. Introduction

Chronic pain of neuropathic origin affects about 8% of the world population and accounts for up to 25% of the total amount of patients with chronic pain. In the clinical setting, patients with neuropathic pain present a range of debilitating symptoms, including spontaneous and evoked pain (e.g., hypersensitivity and hyperalgesia) [1]. Current treatments for neuropathic pain (e.g., antidepressants, anticonvulsants) involve multiple side effects that impair continuation of treatment. Therefore, research identifying safer and more tolerable treatments is needed [2]. Cannabidiol (CBD) is one of the major phytocannabinoids present in the *Cannabis* plant and it was first isolated in 1940 [3]. Due to its favorable safety profile, therapeutic use of CBD has been advocated in treatment of several pathological conditions, including chronic pain [4].

In a model of chronic constriction injury (CCI) of the sciatic nerve, CBD reduced behavioral hypersensitivities to mechanical and heat stimulation after acute [5] and chronic administration [6]. CBD also reduced mechanical hypersensitivity in the model of streptozotocin-induced diabetic neuropathic pain in both mice [7] and rats [8], and attenuated both cold and mechanical hypersensitivity in models of chemotherapy induced neuropathic pain [9,10]. Efficacy of CBD in reducing chronic pain has been reported in multiple models, however, the pharmacological mechanisms through which CBD exerts its antinociceptive effects remain undefined. CBD can activate up to 76 different targets among ion channels, transporters, G-protein coupled receptors (GPCRs) and enzymes [11], in vitro, suggesting that complex polypharmacology underlies the therapeutic effects of CBD.

CBD shows little or no affinity for orthosteric sites at cannabinoid receptors, acting as an allosteric modulator at CB1 receptors and only as a partial agonist at CB2 receptors [12], multiple pre-clinical studies have demonstrated that the analgesic effects of CBD do not involve activation of CB1 or CB2 receptors [8,10,13]. In addition, CBD produced therapeutic effects in a preclinical epilepsy model through antagonism of another GPCR, GPR55, which is highly expressed in the brain [14]. In addition, blockade of GPR55 in the periaqueductal gray matter (PAG), a neural substrate implicated in the descending control of pain, reduces neuropathic nociception in rats [15]. CBD may stimulate *N*-acylphosphatidylethanolamine-specific phospholipase D (NAPE-PLD), an enzyme involved in the production of N-acyl ethanolamines such as anandamide and palmitoylethanolamine, which themselves engage a variety of distinct targets [16]. CBD altered several bioactive lipids in wild-type (WT) mice, but many of these changes were absent in NAPE-PLD knockout mice (NAPE-PLD KO) [16].

We administered CBD acutely and chronically, using both prophylactic and therapeutic dosing schedules, to evaluate the antinociceptive effects of CBD on behavioral hypersensitivities to mechanical and cold stimulation in a model of chemotherapy-induced neuropathic pain (CINP) induced by the taxane paclitaxel. Pharmacological specificity was assessed with selective antagonists of peroxisome proliferator–activated receptors gamma and alpha, as well as cannabinoid receptors type 1 and type 2. To ascertain whether the antinociceptive effects of CBD were dependent upon the enzyme NAPE-PLD, we used a NAPE-PLD knockout (KO) mouse line (Luquet). We also used GPR55 KO mice, to evaluate whether GPR55, implicated in an epilepsy model, could contribute to antinociceptive efficacy of CBD in a model of CIPN. Our studies provide compelling evidence that the antinociceptive effects of CBD require the enzyme NAPE-PLD.

## 2. Methods

### 2.1 Subjects

Age-matched NAPE-PLD KO and WT mice of both sexes, as well as GPR55 KO and WT male mice, were bred at Indiana University. Mice from our breeding colonies were supplemented with age-matched mice, on a C57BL/6 background strain that were purchased from the Jackson laboratories. Animals were kept in standard conditions of temperature (73 ± 2° F), humidity (45%) and under a light/dark cycle (12h/12h). All the experimental procedures were approved by Bloomington Institutional Animal Care and Use Committee of Indiana University and followed the guidelines for the treatment of animals of the International Association for the Study of Pain [17].

### 2.2 Drugs and Chemicals

Paclitaxel (Tecoland Coporation, Irvine, CA) was dissolved in a vehicle consisting of cremophor (Sigma-Aldrich, St. Louis, MO), ethanol (Sigma-Aldrich), and saline (Aquilite System, Hospira Inc, Lake Forest, IL) at a ratio of 1:1:18 as previously described [18–21]. CBD obtained from the NIDA Drug Supply Program was supplemented with CBD provided by Joshua Harnett (Blue Sky Processing LLC, Seabrook, SC), with 99% purity. CBD was prepared in a vehicle composed of dimethysulfoxide (Sigma-Aldrich, St. Louis, MO), ethanol, emulphor (Alkamuls EL 620L, Solvay, Princeton, NJ) and saline at a ratio of 5:2:2:16. AM251, AM630, GW6471 and GW9662 (Cayman, MI, USA) were diluted in a vehicle consisting of DMSO, ethanol, cremophor and saline at a 1:1:1:17 ratio[22,23]. Paclitaxel was given in a volume of 6.67 ml/kg. All receptor antagonists and CBD were given in an injection volume of 5 ml/kg.

### 2.3 Paclitaxel-induced neuropathic pain

Paclitaxel (4 mg/kg/day, i.p.) or its cremophor-based vehicle was administered once daily on alternate days for a total of 4 injections. Responsiveness to mechanical and cold sensitivity were assessed as a baseline (i.e., pre-injection response) and every 3-5 days after the initiation of paclitaxel dosing in mice for up to 28 days.

### 2.4 Assessment of mechanical paw withdrawal thresholds

Paw withdrawal thresholds (grams) and development of mechanical hypersensitivity were evaluated as previously described by our groups [18–21]. Briefly, animals were habituated for 30 minutes in individual inverted plexiglass chambers placed on a metal mesh platform positioned on a stable wooden table. Paw withdrawal thresholds were evaluated with a semi-flexible 90-g probe connected to an electronic von Frey anesthesiometer (IITC Life Science, Woodland Hills, CA). Mechanical paw withdrawal threshold was defined as the grams necessary to produce paw withdrawal and was recorded in duplicate for each hind paw, with a 2-minute interval between stimulations to avoid sensitization. Mechanical withdrawal threshold was then averaged across paws to provide a single measurement per animal per timepoint because our previous work has shown that the present paclitaxel dosing paradigm produces bilateral neuropathy that does not differ between paws.

### 2.5 Assessment of cold hypersensitivity

Hypersensitivity to cold was evaluated immediately after the assessment of mechanical paw withdrawal thresholds in the same animals as described previously [18,21]. A 1 mL syringe with needle removed was filled with acetone (Sigma-Aldrich). The plunger of the syringe was pushed gently to form an acetone drop at the tip, which was then applied to the plantar surface of the hind paw with care taken to avoid contacting the paw with the syringe tip and applying mechanical stimulation. The time in seconds that animal attended on the stimulated paw (i.e., elevating, biting, licking, shaking, or flinching) was measured in triplicate for each paw with a few-minute interval between stimulations to avoid sensitization. The duration of response to cold stimulation was then averaged across paws.

### 2.6 Experimental procedures

The therapeutic effect of CBD was evaluated acutely, as well as during the development and maintenance phases of chemotherapy-induced peripheral neuropathy induced by paclitaxel. The following experimental procedures were employed:

Experiment 1: We examined the impact of acute treatment with CBD in male mice with established paclitaxel-induced hypersensitivity using a within subject dosing paradigm. Mice received injections of CBD (1, 3, 10 or 30 mg/kg, i.p.) or vehicle starting on day 16 after the first injection of paclitaxel (4 mg/kg, i.p.). Increasing doses were given in an interval of 2-3 days (**Figure 1A**). Mechanical threshold and response to acetone were reevaluated 30 minutes after treatment.

**Figure 1.**
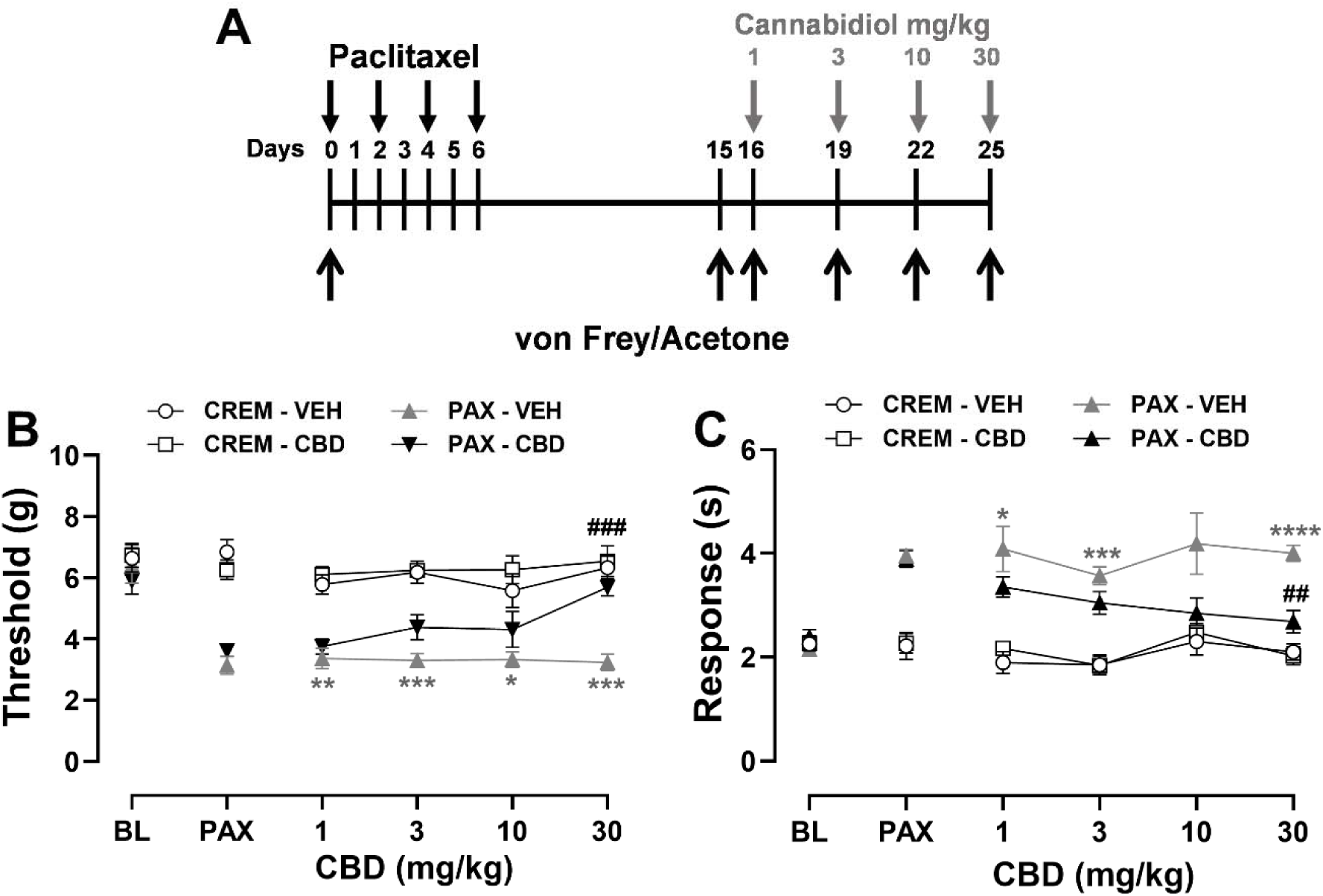
Cannabidiol (CBD) acutely attenuates paclitaxel-induced mechanical and cold hypersensitivity in a dose-dependent manner. Dose response curve of CBD (1-30 mg/kg, i.p.) in suppressing paclitaxel-induced hypersensitivity to (**B**) mechanical and (**C**) cold stimulation in male mice (n=6). Data are expressed as mean ± SEM. Two-way ANOVA followed by Bonferroni’s *post hoc* test. *\*\*\*\*p<0.0001, ***p<0.001, **p<0.01, *p<0.05 CREM VEH vs. PAX-Veh . ###p<0.001, ##p<0.01 PAX-Veh vs. PAX-CBD*.

**Figure 2.**
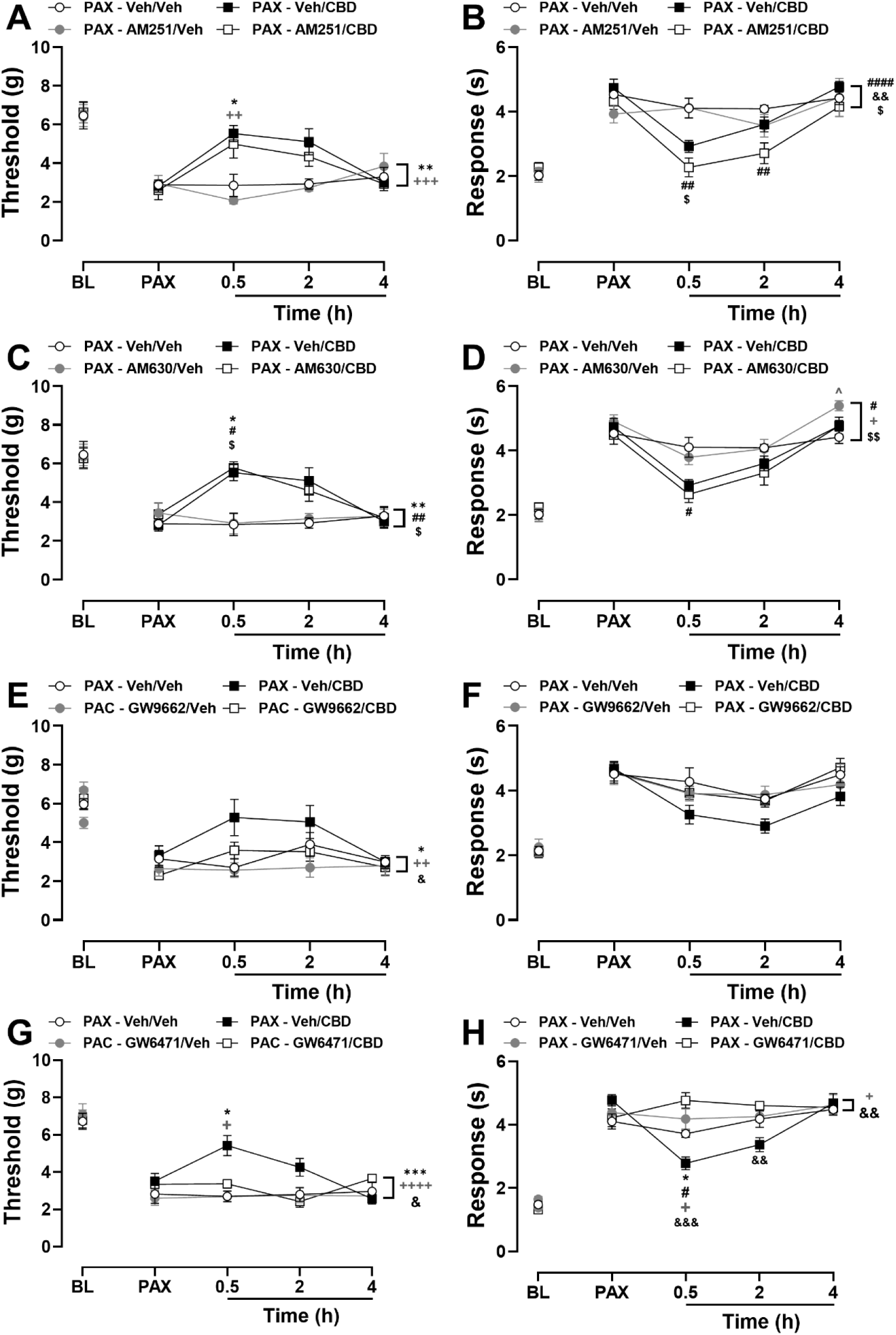
PPAR receptor antagonists, but not cannabinoid receptor antagonists attenuate the antinociceptive effect of CBD (30 mg/kg, i.p.) on paclitaxel-induced hypersensitivity in mice. CB1 receptor antagonist AM251 (5mg/kg, i.p), does not attenuate the antinociceptive effect of CBD on paclitaxel-induced (**A**) mechanical (**B**) cold hypersensitivity. CB2 receptor antagonist, AM630 (5mg/kg, i.p), does not attenuate the antinociceptive effect of CBD on paclitaxel-induced (**C**) mechanical (**D**) cold hypersensitivity. PPARγ receptor antagonist, GW9662 (2 mg/kg, i.p), attenuates the antinociceptive effect of CBD on paclitaxel-induced (**E**) mechanical hypersensitivity, but not (**F**) cold hypersensitivity. PPARα receptor antagonist, GW6471 (4 mg/kg, i.p), attenuates the antinociceptive effect of CBD on paclitaxel-induced (**G**) mechanical and (**H**) cold hypersensitivity. Data are expressed as mean ± SEM. Two-way ANOVA followed by Bonferroni’s *post hoc* test. Interaction between time and treatment: *\*p<0.05 PAX-Veh/CBD vs. PAX-Veh/Veh; ++p<0.01, +p<0.05 PAX-Veh/CBD vs. PAX-Antagonist/Veh; ##p<0.01, #p<0.05 PAX-Veh/Veh vs. PAX-Antagonist/CBD; $p<0.05 PAX-Antagonist/Veh vs. PAX-Antagonist/CBD; &&&p<0.001, &&p<0.01 PAX-Veh/CBD vs. PAX-Antagonist/CBD; ^p<0.05 PAX-Veh/Veh vs. PAX-Antagonist/Veh. Brackets show main treatment effect: ***p<0.001, **p<0.01, *p<0.05 PAX-Veh/CBD vs. PAX-Veh/Veh; ++++p<0.0001, ++p<0.01, +p<0.05 PAX-Veh/CBD vs. PAX-Antagonist/Veh; ####p<0.01, ##p<0.01, #p<0.05 PAX-Veh/Veh vs. PAX-Antagonist/CBD; $p<0.01, $p<0.05 PAX-Antagonist/Veh vs. PAX-Antagonist/CBD; &&p<0.01, &p<0.05 PAX-Veh/CBD vs. PAX-Antagonist/CBD*.

Experiment 2: We assessed the impact of pharmacological antagonists of diverse receptor targets linked to mechanism of action of CBD, including: 1) CB1 receptor antagonist (AM251, 5 mg/kg, i.p.); 2) CB2 receptor antagonist (AM630, 5 mg/kg, i.p.), 3) PPARα antagonist (GW6471, 4 mg/kg, i.p.); or 4) PPARγ antagonist (GW9662, 2 mg/kg, i.p.). Antagonists were administered 30 minutes before an acute CBD injection (30 mg/kg, i.p.) during the maintenance phase of neuropathic nociception in paclitaxel-treated mice tested 2-3 weeks after initial paclitaxel dose. Animals were tested for mechanical paw withdrawal thresholds 0.5, 2 and 4 hours after CBD injection. Antagonist treatment was given in a within subject manner in different groups, with washout between successive antagonist treatments: group 1: AM251, AM630 and GW9662; group 2: GW6471. Treatments were administered with a 2–4-day interval between each antagonist. Mechanical paw withdrawal thresholds were reassessed to ensure lack of any residual effect was absent before advancing to the next treatment. Groups were randomized before each treatment to assure a consistent drug/vehicle injection history.

Experiment 3: We assessed CBD treatment during the maintenance phase of paclitaxel-induced hypersensitivity in NAPE-PLD KO mice and their respective WT aged-matched groups. Mice of both sexes received 7 daily injections of CBD (10 mg/kg/day, i.p.) or vehicle starting 15 days after the first injection of paclitaxel (4 mg/kg, i.p) (**Figure 3A and 4A**).

**Figure 3.**
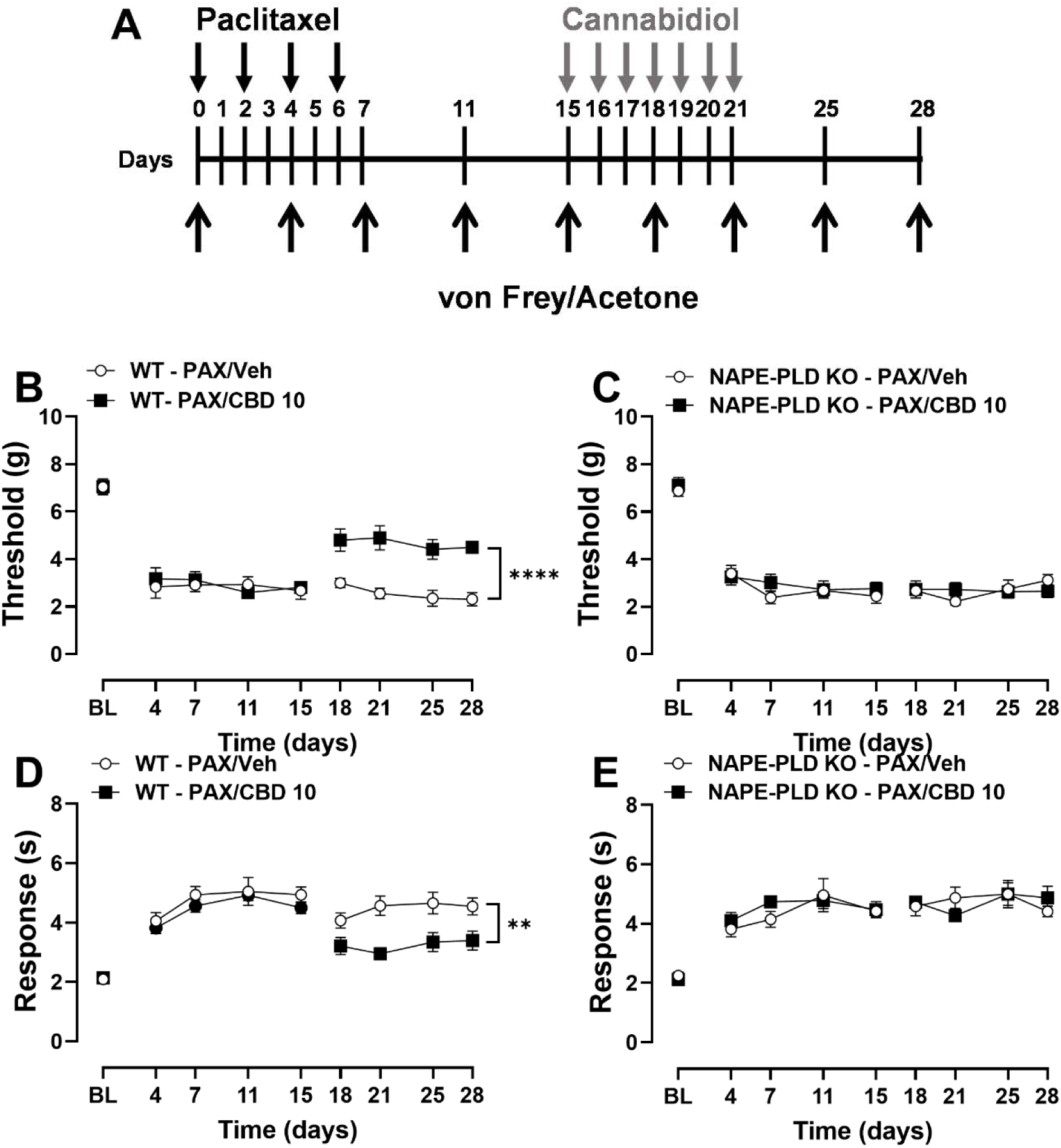
CBD reduces the maintenance of paclitaxel-induced hypersensitivity in a NAPE-PLD dependent manner in male mice. Treatment with CBD (10 mg/kg, i.p.; 7 daily injections) given during the maintenance phase of paclitaxel-induced neuropathic pain attenuated mechanical hypersensitivity in (**B**) WT, but not in (**C**) NAPE-PLD knock out (KO) male mice. Similarly, CBD attenuated paclitaxel-induced cold hypersensitivity in (D) WT, but not (E) NAPE-PLD KO male mice. Data are expressed as mean ± SEM. Two-way ANOVA followed by Bonferroni’s *post hoc* test. *Brackets show main treatment effect: ****p<0.0001, **p<0.01 WT – PAX/Veh vs. WT – PAX/CBD 10*.

**Figure 4.**
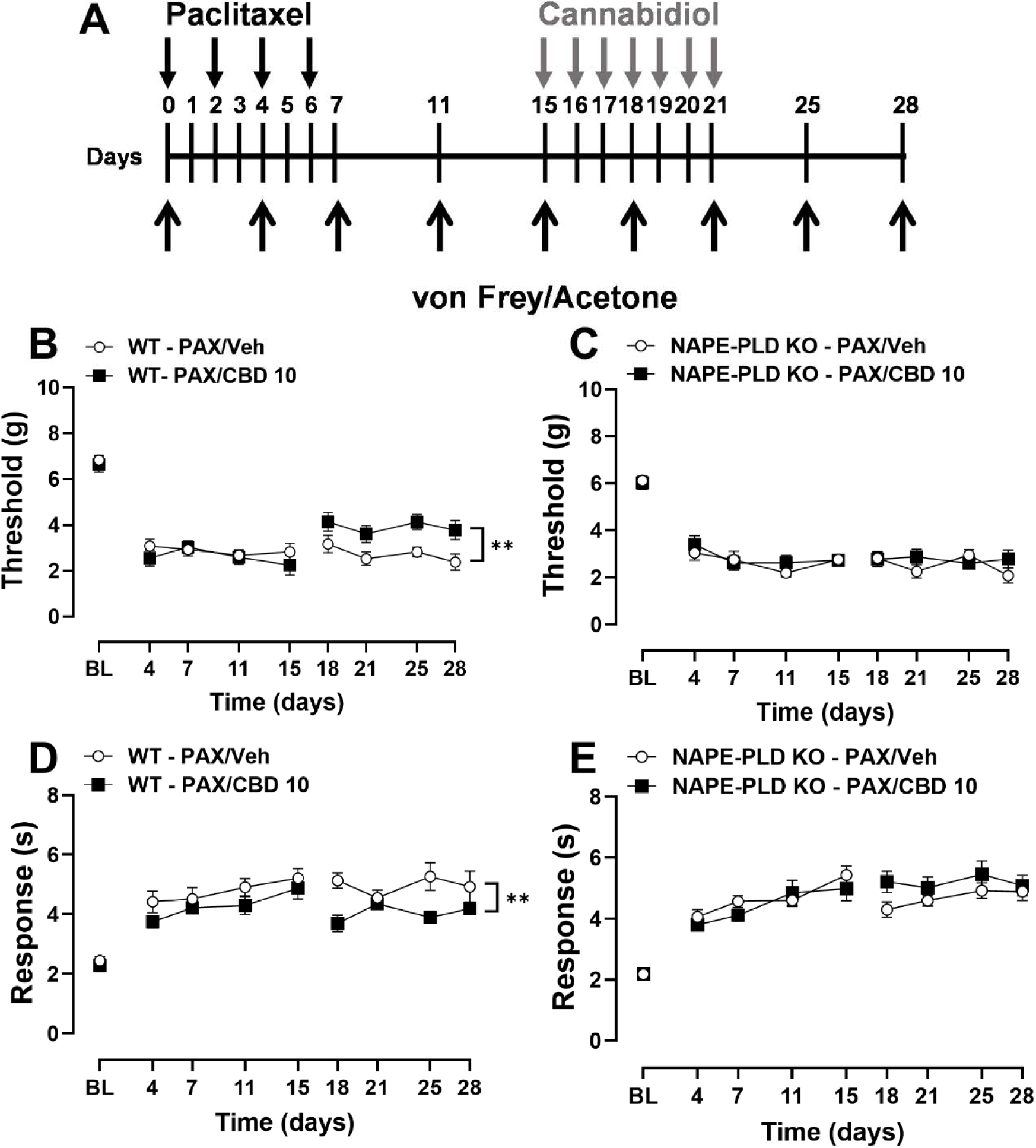
CBD reduces the maintenance of paclitaxel-induced hypersensitivity in a NAPE-PLD dependent manner in female mice. Treatment with CBD (10 mg/kg, i.p.; 7 daily injections) given during the maintenance phase of paclitaxel-induced neuropathic pain attenuated mechanical hypersensitivity in (**B**) WT, but not in (**C**) NAPE-PLD KO female mice. Similarly, CBD attenuated paclitaxel-induced cold hypersensitivity in (**D**) WT, but not (**E**) NAPE-PLD KO female mice. Data are expressed as mean ± SEM. Two-way ANOVA followed by Bonferroni’s *post hoc* test. *Brackets show main treatment effect: **p<0.01 WT – PAX/Veh vs. WT – PAX/CBD 10*.

Experiment 4: We assessed CBD treatment during the maintenance phase of paclitaxel-induced hypersensitivity in GPR55 KO mice and their respective WT aged-matched groups. WT and GPR55 KO male mice received 7 daily injections of CBD (10 mg/kg/day, i.p.) or vehicle starting 15 days after the first injection of paclitaxel (4 mg/kg, i.p) (**Figure 5A**).

**Figure 5.**
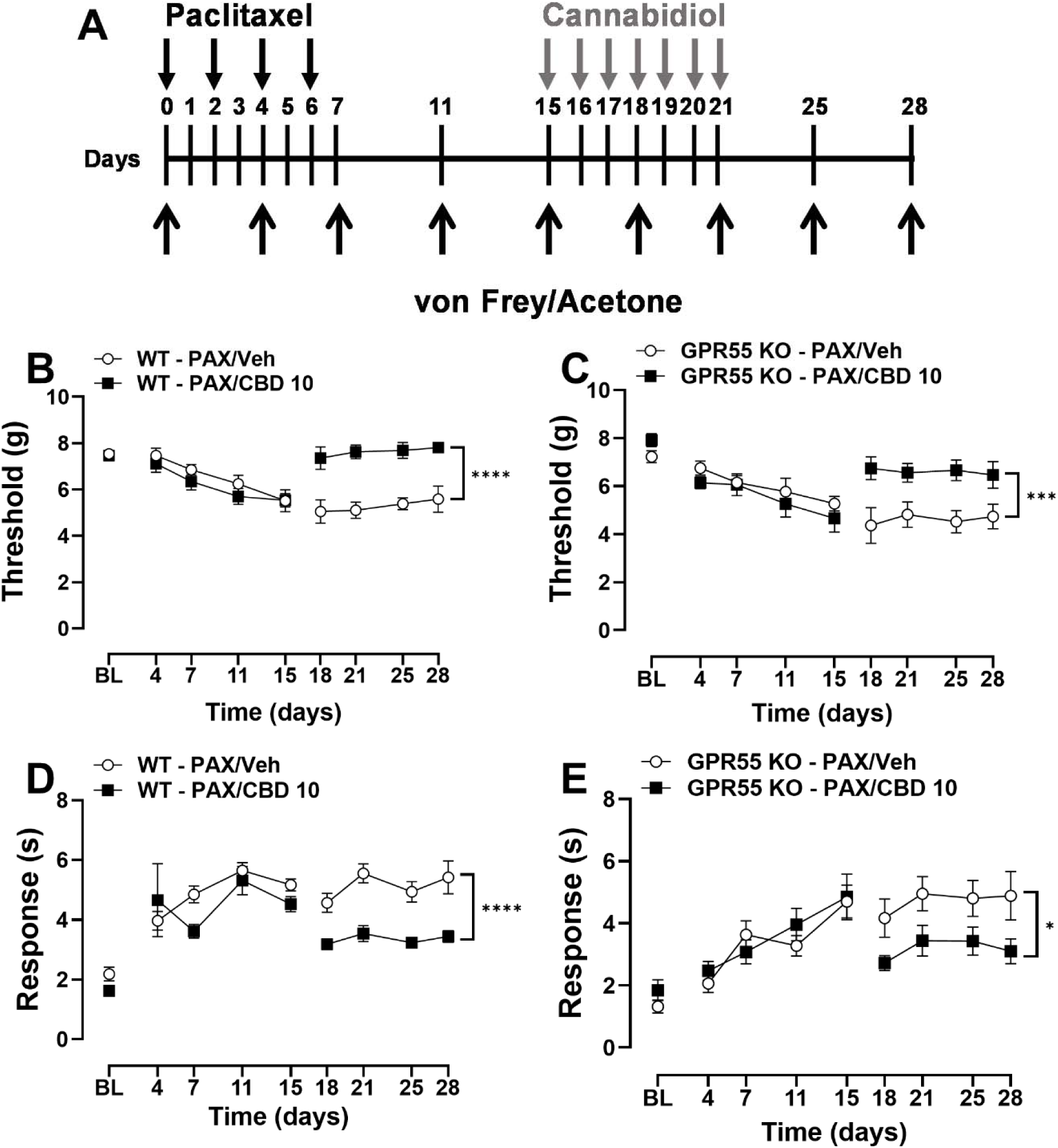
CBD reduces the maintenance of paclitaxel-induced hypersensitivity in both WT and GPR55 receptor KO mice. Treatment with CBD (10 mg/kg, i.p.; 7 daily injections) given during the maintenance phase of paclitaxel-induced neuropathic pain attenuated mechanical hypersensitivity in (**B**) WT and (**C**) GPR55 KO male mice. Similarly, CBD attenuated paclitaxel-induced cold hypersensitivity in (**D**) WT and (**E**) GPR55 KO male mice. Data are expressed as mean ± SEM. Two-way ANOVA followed by Bonferroni’s *post hoc* test. *Brackets show main treatment effect: ****p<0.0001, ***p<0.001, *p<0.05 WT – PAX/Veh vs. WT – PAX/CBD 10 or GPR55 KO – PAX/Veh vs. GPR55 – PAX/CBD 10*.

Experiment 5: We assessed the impact of prophylactic CBD treatment during the development phase of paclitaxel-induced hypersensitivity in NAPE-PLD KO and WT aged-matched groups. Mice of both sexes received 7 daily injections of CBD (10 mg/kg/day, i.p.) or vehicle starting 1 hour before the first injection of paclitaxel (4 mg/kg, i.p) (**Figure Figure 6A and 7A**).

**Figure 6.**
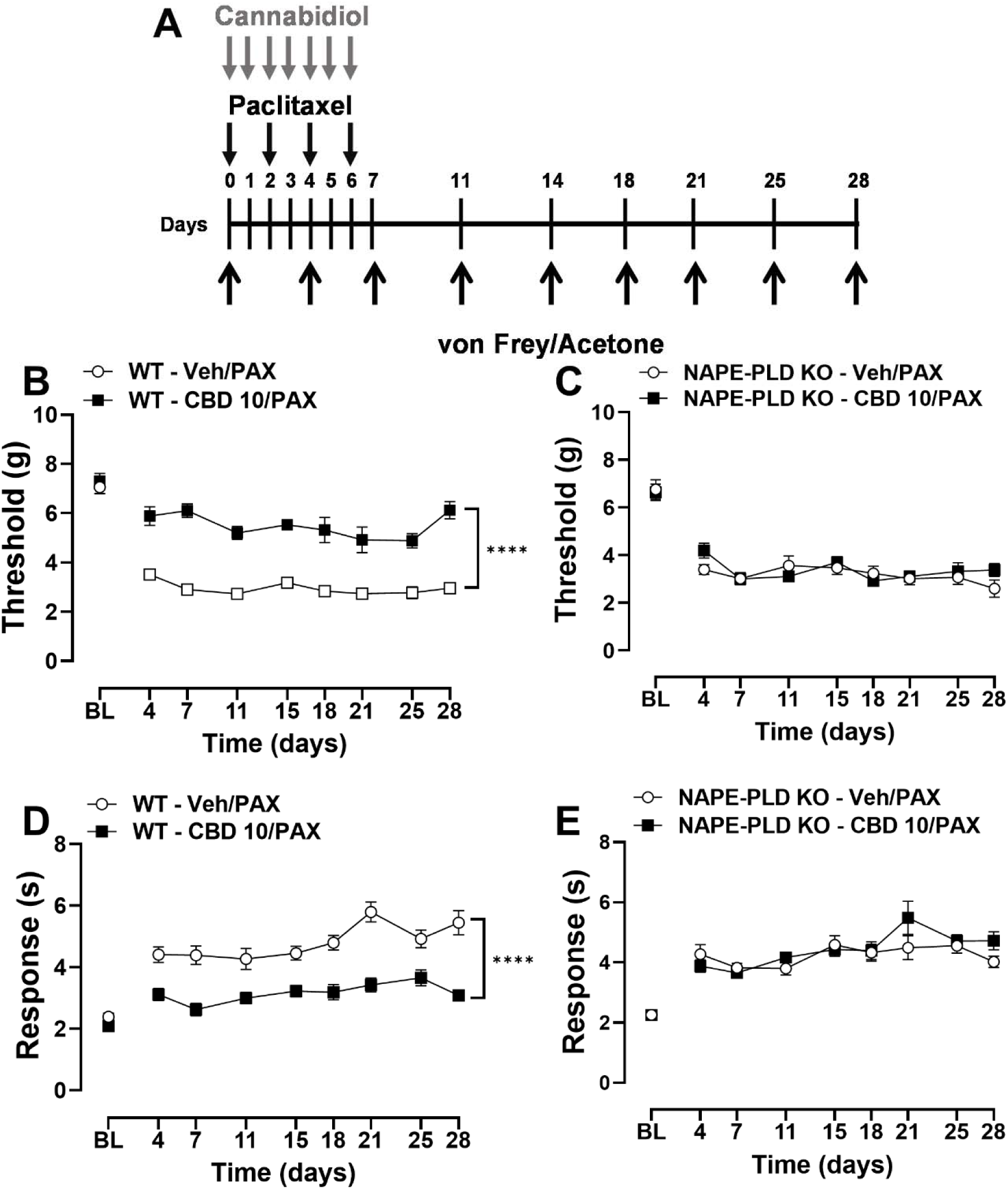
CBD attenuates the development of paclitaxel-induced hypersensitivity in a NAPE-PLD dependent manner in male mice. Treatment with CBD (10 mg/kg, i.p.; 7 daily injections) given during the development phase of paclitaxel-induced neuropathic pain attenuated mechanical hypersensitivity in (**B**) WT, but not (**C**) NAPE-PLD KO male mice. CBD attenuated paclitaxel-induced cold hypersensitivity in (**D**) WT, but not in (**E**) NAPE-PLD KO male mice. Data are expressed as mean ± SEM. Two-way ANOVA followed by Bonferroni’s *post hoc* test. *Brackets show main treatment effect: ****p<0.0001 WT – Veh/PAX vs. WT – CBD 10/PAX*.

**Figure 7.**
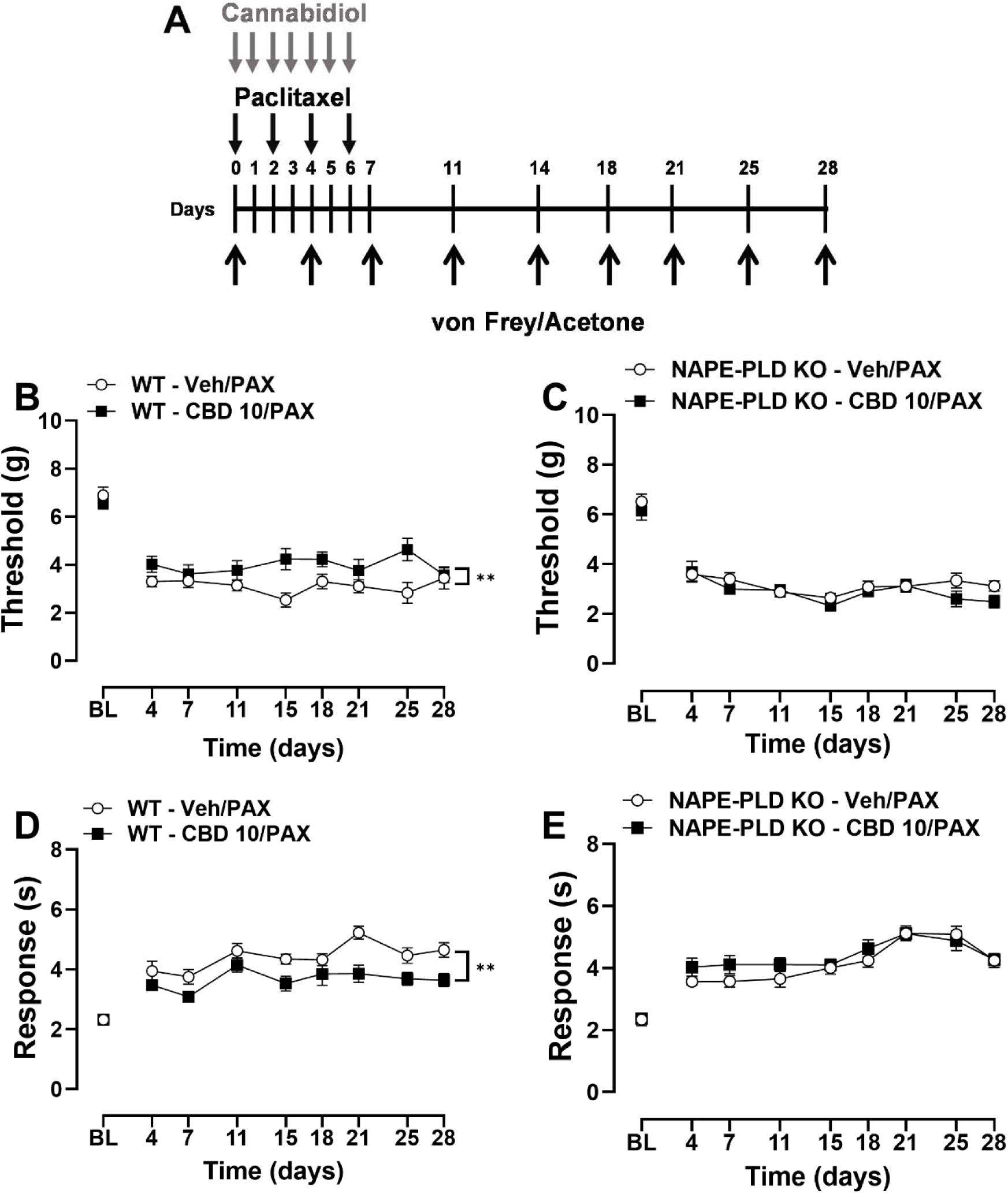
CBD attenuates the development of paclitaxel-induced hypersensitivity in a NAPE-PLD dependent manner in female mice. Treatment with CBD (10 mg/kg, i.p.; 7 daily injections) given during the development phase of paclitaxel-induced neuropathic pain attenuated mechanical hypersensitivity in (**B**) WT, but not (**C**) NAPE-PLD KO female mice. CBD attenuated paclitaxel-induced cold hypersensitivity in (**D**) WT, but not in (**E**) NAPE-PLD KO female mice. Data are expressed as mean ± SEM. Two-way ANOVA followed by Bonferroni’s *post hoc* test. *Brackets show main treatment effect: **p<0.0001 WT – Veh/PAX vs. WT – CBD 10/PAX*.

Experiment 6: We evaluated the impact of prophylactic treatment with CBD or its vehicle during the development phase of paclitaxel-induced hypersensitivity using WT and GPR55 KO male mice and their respective WT aged-matched groups. Mice received 7 daily injections of CBD (10 mg/kg/day, i.p.) or vehicle starting 1 hour before the first injection of paclitaxel (4 mg/kg, i.p) (**Figure 8A**).

**Figure 8.**
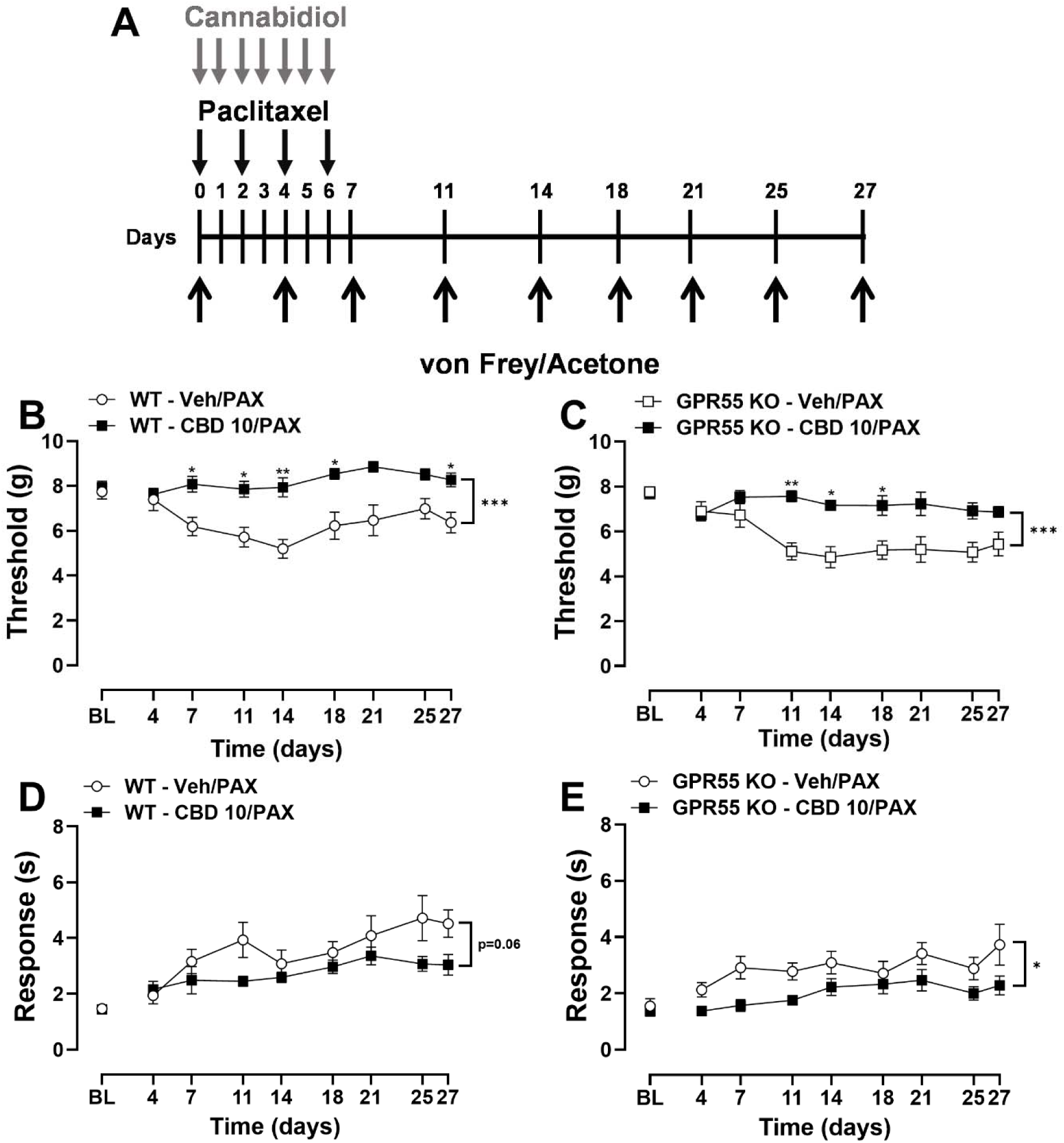
CBD attenuates the development of paclitaxel-induced hypersensitivity in both WT and GPR55 receptor KO mice. Treatment with CBD (10 mg/kg, i.p.; 7 daily injections) given during the development phase of paclitaxel-induced neuropathic pain attenuated mechanical hypersensitivity in (**B**) WT and (**C**) GPR55 KO male mice. Similarly, CBD attenuated paclitaxel-induced cold hypersensitivity in (**D**) WT and (**E**) GPR55 KO male mice. Data are expressed as mean ± SEM. Two-way ANOVA followed by Bonferroni’s *post hoc* test. Interaction between time and treatment: *\*\*p<0.01, *p<0.05 WT – Veh/PAX vs. WT – CBD 10/PAX or GPR55 KO – Veh/PAX vs. GPR55 KO – CBD 10/PAX. Brackets show main treatment effect: ***p<0.001, *p<0.05 WT – Veh/PAX vs. WT – CBD 10/PAX or GPR55 KO – Veh/PAX vs. GPR55 KO – CBD 10/PAX*.

### 2.8 Statistical analysis

Two-way repeated measures analysis of variance (ANOVA) was employed for the analyses of main effects of treatment, time, and interaction between treatment and time. Bonferroni post hoc tests were used for pairwise multiple comparisons. Statistical analyses and graph design were performed using GraphPad Prism version 9.4.0. Data were expressed as mean ± S.E.M. and *p* < 0.05 was considered statistically significant.

## 3. Results

### 3.1 Acute treatment with CBD attenuated paclitaxel-induced mechanical and cold hypersensitivity

Treatment with paclitaxel (4 mg/kg, i.p. x 4 injections) induced mechanical (**Figure 1B**) and cold hypersensitivity (**Figure 1C**) that was well-established prior to pharmacological manipulations (i.e., initiated 16 days after the first injection). Paclitaxel produced robust mechanical hypersensitivity (vs Cremophor-treated mice, **Figure 1B**). CBD (30 mg/kg, i.p.), given in a within subject paradigm, alleviated paclitaxel-induced mechanical hypersensitivity in mice (**Figure 1B**: Treatment: F_3, 20_ = 46.68, p<0.0001; Dose: F_2.794, 55.89_=2.903, p=0.0462; Interaction: F_8.383, 55.89_ = 1.208, p=0.3101), when compared to vehicle (p=0.0005), and normalized responses to those observed in groups receiving Cremophor-vehicle in lieu of paclitaxel (p>0.99). Paclitaxel treatment also produced cold hypersensitivity (vs Cremophor-treated mice, **Figure 1C**). CBD (30 mg/kg, i.p.) alleviated paclitaxel-induced cold hypersensitivity (**Figure 1C**: Treatment: F_3, 20_ = 26.58, p<0.0001; Dose: F_2.557, 51.14_=2.265, p=0.1012; Interaction: F_7.671, 51.14_ = 1.030, p=0.4246), compared to vehicle-treated mice (p=0.0049). No alterations in response to cold stimulation were observed in the Cremophor group treated with CBD (p=0.2381).

### 3.2 Acute antiallodynic effect of CBD is suppressed by PPAR receptor antagonists, but not cannabinoid receptor antagonists

In general, our paclitaxel dosing regimen reduced mechanical paw withdrawal threshold and responsiveness to acetone, consistent with development of mechanical and cold hypersensitivity, respectively (**Figure 2A-H**, BL vs PAX). Acute treatment with CBD (30 mg/kg, i.p.) attenuated mechanical and/or cold hypersensitivity 30 minutes after injection (**Figure 2A-H**, PAC vs Post treatment time course; For treatment effect statistics see **Table 1**).

**Table 1.**

| Figure | Treatment | Time | Interaction |
| --- | --- | --- | --- |
| 2A | F (3, 18) = 6.802, P=0.0029 | F (2.805, 50.50) = 3.984, P=0.0144 | F (8.416, 50.50) = 3.834, P=0.0012 |
| 2B | F (3, 18) = 10.93, P=0.0003 | F (2.562, 46.11) = 17.11, P<0.0001 | F (7.686, 46.11) = 3.117, P=0.0073 |
| 2C | F (3, 18) = 7.456, P=0.0019 | F (2.736, 49.25) = 5.666, P=0.0027 | F (8.208, 49.25) = 3.407, P=0.0032 |
| 2D | F (3, 18) = 3.817, P=0.0281 | F (2.706, 48.71) = 40.79, P<0.0001 | F (8.119, 48.71) = 3.093, P=0.0065 |
| 2E | F (3, 18) = 3.793, P=0.0287 | F (2.522, 45.39) = 3.639, P=0.0254 | F (7.565, 45.39) = 1.599, P=0.1552 |
| 2F | F (3, 18) = 3.068, P=0.0543 | F (2.712, 48.81) = 10.64, P<0.0001 | F (8.135, 48.81) = 1.010, P=0.4415 |
| 2G | F (3, 20) = 9.204, P=0.0005 | F (2.811, 56.22) = 1.887, P=0.1457 | F (8.432, 56.22) = 3.524, P=0.0020 |
| 2H | F (3, 20) = 4.302, P=0.0170 | F (2.788, 55.75) = 7.334, P=0.0004 | F (8.363, 55.75) = 4.720, P=0.0002 |

Administration of CB1 receptor antagonist AM251 (5 mg/kg, i.p.), prior to CBD treatment, did not reliably block the antinociceptive effect of CBD over mechanical hypersensitivity throughout the course of the study (**Figure 2A**; p>0.99). CBD treatment attenuated mechanical hypersensitivity 30 minutes post-treatment (vs. vehicle-vehicle or AM251-vehicle, p<0.0266; **Figure 2A**). Although AM251-CBD differed from vehicle-CBD treatment in cold hypersensitivity (p<0.0039), it still significantly reduced cold hypersensitivity compared with vehicle–vehicle and AM251-vehicle treatments (p<0.0117; **Figure 2B**). AM251-CBD attenuated cold hypersensitivity at 30 minutes and 2 hours versus vehicle-treated mice (p<0.0403), and at 30 minutes versus AM251-vehicle-treated mice (p=0.0171; **Figure 2B**).

CBD treatment attenuated mechanical hypersensitivity in paclitaxel-treated mice (vs. vehicle-vehicle, p=0.0038), an effect independent of pre-injection with the CB2 receptor antagonist AM630 (vs. AM630-CBD, p=0.0011; **Figure 2C**). AM630-CBD combination attenuated mechanical hypersensitivity across the entire course of the study (vs. AM630-vehicle, p=0.0258). CBD alleviated mechanical hypersensitivity at 30 minutes post-treatment compared with the vehicle–vehicle (p=0.0266) and AM630-vehicle groups (p=0.0377), regardless of AM630 pre-injection (vehicle-vehicle vs. AM630-CBD, p=0.0139; **Figure 2C**).

CBD attenuated cold hypersensitivity when compared to AM630-vehicle group (p=0.0344). AM630 did not block the effects of CBD on cold hypersensitivity (vs. vehicle-vehicle and AM630-vehicle groups, p<0.0279; **Figure 2D**). AM630-CBD attenuated cold hypersensitivity at 30 minutes post-treatment (vs. vehicle-vehicle, p=0.0283). Although a significant effect of treatment was detected at 240 minutes post injection in mice given AM630 only (AM630-vehicle vs vehicle-vehicle, p=0.0304), no difference from baseline response to cold stimulus after paclitaxel injection was detected (p=0.4725).

Combination of PPARγ antagonist GW9662 and CBD attenuated paclitaxel induced mechanical hypersensitivity overall (Treatment: F_3, 18_ = 3.793, p=0.0287; **Figure 2E**), but the interaction was not significant (F_7.565, 45.39_ = 1.599, p=0.1552; **Figure 2E**). CBD attenuated paclitaxel-induced mechanical hypersensitivity, when compared to the groups of mice treated with only vehicle (p=0.0498) or GW9662 alone (p=0.0026), and GW9662 pre-treatment blocked the overall antiallodynic effect of CBD alone (p=0.0147; **Figure 2E**). In this subset of data, no significant effects of treatment were detected for CBD over paclitaxel-induced cold hypersensitivity (Treatment: F_3, 18_ = 3.068, p=0.0543; **Figure 2F**).

Moreover, CBD attenuated paclitaxel-induced mechanical hypersensitivity compared to all other groups and PPARα antagonist administration blocked the effect of CBD (p<0.0377 vehicle-CBD vs. all other groups, **Figure 2G**). CBD attenuated mechanical hypersensitivity at 30 minutes pos-treatment (vs. vehicle-vehicle or GW6471-vehicle, p<0.0153; **Figure 2G**). Accordingly, CBD attenuated cold hypersensitivity in comparison to GW6471-vehicle group (p=0.0494; **Figure 2H**). GW6471 pre-treatment blocked the anti-allodynic effect of CBD over cold hypersensitivity in paclitaxel-treated mice (p=0.0022). Co-administration of GW6471 and CBD led to an exacerbated response to acetone in mice, compared to vehicle-vehicle group at 30 minutes (p=0.0364). Vehicle-CBD treatment attenuated cold hypersensitivity at 30 minutes post-treatment relative to all groups (p<0.0338). Pre-treatment with GW6471 significantly blunted the CBD-induced attenuation of cold hypersensitivity at 30 minutes and 2 hours post treatment (vs vehicle-CBD, p<0.0085; **Figure 2H**).

### 3.3 Treatment with CBD reduced the maintenance of paclitaxel-induced mechanical and cold hypersensitivity in a NAPE-PLD dependent manner in male and female mice

Overall, paw withdrawal thresholds and response to acetone in WT and NAPE-PLD KO male mice were decreased during the development phase of paclitaxel-induced peripheral neuropathy (BL to day 15, **Figures 3B-E**). Overall, in WT male mice (**Figure 3B**), daily administration of a subthreshold dose of CBD (10 mg/kg, i.p.; day 15 to 21) increased mechanical paw withdrawal thresholds during the maintenance phase of paclitaxel-induced mechanical hypersensitivity (**Figure 3B**: Treatment: F_1, 14_ = 78.74, p<0.0001; Time: F_2.205, 30.87_=1.002, p=0.3781; Interaction: F_3, 42_ = 0.2080, p=0.8903). Daily administration of CBD (10 mg/kg, i.p.) did not increase mechanical paw withdrawal thresholds in NAPE-PLD KO male mice during the maintenance phase of paclitaxel-induced mechanical hypersensitivity (**Figure 3C**: Treatment: F_1, 17_ = 0.0004, p=0.3196; Time: F_2.728, 46.38_=0.7147, p=0.5357; Interaction: F_3, 51_ = 1.005, p=0.3984).

In WT male mice, daily injection of CBD (10 mg/kg, i.p.; day 15 to 21) decreased response to acetone overall (**Figure 3D**: Treatment: F_1, 14_ = 13.18, p=0.0027; Time: F_2.071, 28.99_=1.410, p=0.2606; Interaction: F_3, 42_ = 1.194, p=0.3236). Daily injection with CBD did not alter responses to acetone in NAPE-PLD KO mice, when in comparison to vehicle-treated groups (**Figure 3E**: Treatment: F_1, 17_ = 2.044e-005, p=0.9964; Time: F_2.447, 41.59_=1.254, p=0.300; Interaction: F_3, 51_ = 1.654, p=0.1885).

Paclitaxel treatment decreased paw withdrawal thresholds and response to acetone in female mice in WT or NAPE-PLD KO conditions (BL to day 15, **Figures 4B-E**). In WT female mice, CBD (10 mg/kg, daily, 7 days) given during the maintenance phase of paclitaxel-induced neuropathy increased mechanical paw withdrawal thresholds (**Figure 4B**: Treatment: F_1, 14_ = 12.00, p=0.0038; Time: F_2.510, 35.14_=2.039, p=0.1352; Interaction: F_2.510, 35.14_ = 0.2312, p=0.8412). In contrast, CBD treatment did not alter the mechanical thresholds of NAPE-PLD KO female mice in the maintenance phase of paclitaxel-induced hypersensitivity (**Figure 4C**: Treatment: F_1, 16_ = 0.8027, p=0.3836; Time: F_2.344, 37.50_ = 0.9018, p=0.4285; Interaction: F_2.344, 35.50_ = 1.901, p=0.1574).

CBD treatment also attenuated responses to acetone in WT female mice during the maintenance phase of paclitaxel-induced neuropathy overall (**Figure 4D**: Treatment: F_1, 14_ = 8.684, p=0.0106; Time: F_2.409, 33.72_ = 0.1853, p=0.8681; Interaction: F_2.409, 33.72_ = 2.428, p=0.0942). In contrast, no effect of CBD treatment was detected in the response to acetone of NAPE-PLD KO female mice during the maintenance phase of paclitaxel-induced neuropathy (**Figure 4E**: Treatment: F_1, 16_ = 2.235, p=0.1544; Time: F_2.699, 43.19_ = 1.376, p=0.2637; Interaction: F_2.699, 43.19_ = 0.7966, p=0.4909).

### 3.4 Chronic treatment with CBD reduced the maintenance of paclitaxel-induced mechanical and cold hypersensitivity in both wild-type and GPR55 knock out mice

We also investigated if GPR55 receptors are involved in the anti-allodynic effect of CBD during the maintenance phase of paclitaxel-induced peripheral neuropathy. Overall, male WT and GPR55 KO mice that were subjected to paclitaxel treatment showed a decrease in mechanical paw withdrawal thresholds and increase in response to acetone, indicative of development of mechanical and cold hypersensitivity (BL to day 15, **Figures 5B-E**). Chronic CBD treatment (10 mg/kg, daily, 7 days) given during the maintenance phase of paclitaxel-induced neuropathy increased paw withdrawal thresholds in WT (**Figure 5B**: Treatment: F_1, 14_ = 49.76, p<0.0001; Time: F_2.484, 34.78_=0.6705, p=0.5490; Interaction: F_2.484, 34.78_ = 0.0608, p=0.9654) and GPR55 KO male mice overall (**Figure 5C**: Treatment: F_1, 14_ = 17.13, p=0.0010; Time: F_2.250, 31.49_=0.0316, p=0.9782; Interaction: F_2.250, 31.49_ = 0.2392, p=0.8133). In addition, CBD treatment attenuated cold-hypersensitivity during the maintenance phase of paclitaxel-induced neuropathy in WT (**Figure 5D**: Treatment: F_1, 14_ = 36.85, p<0.0001; Time: F_2.346, 32.84_=2.521, p=0.0878; Interaction: F_2.346, 32.84_ = 0.5588, p=0.6043) and GPR55 (**Figure 5E**: Treatment: F_1, 14_ = 6.079, p=0.0272; Time: F_2.826, 39.57_=1.842, p=0.1582; Interaction: F_2.826, 39.57_ = 0.1312, p=0.9332).

### 3.5 Treatment with CBD attenuates the development of paclitaxel-induced mechanical and cold hypersensitivity in a NAPE-PLD dependent manner in mice

Overall, male mice that received prophylactic treatment with vehicle before paclitaxel developed decreased paw withdrawal thresholds and increased response to acetone (BL-Day 28). CBD treatment (10 mg/kg, daily, 7 days) given before and during paclitaxel regimen attenuated the development of paclitaxel-induced mechanical hypersensitivity in a time dependent manner in WT (**Figure 6B**: Treatment: F_1, 16_ = 251.2, p<0.0001; Time: F_4.509, 72.15_=3.267, p=0.0127; Interaction: F_4.509, 72.15_ = 1.193, p=0.3217), but not in NAPE-PLD KO male mice, when compared to paclitaxel-treated mice given vehicle (**Figure 6C**: Treatment: F_1, 14_ = 1.168, p=0.2980; Time: F_3.938, 55.13_=2.391, p=0.0626; Interaction: F_3.938, 55.13_ = 1.403, p=0.2456). In addition, CBD treatment attenuated the development of cold hypersensitivity in WT paclitaxel-treated mice in a time dependent manner, compared to the WT group of male mice that was given vehicle prophylactically (**Figure 6D**: Treatment: F_1, 16_ = 49.10, p<0.0001; Time: F_4.711, 75.38_=5.341, p=0.0004; Interaction: F_4.711, 75.38_ = 2.213, p=0.0653). This treatment effect was not observed in NAPE-PLD KO mice, only a time effect was detected (**Figure 6E**: Treatment: F_1, 14_ = 0.7186, p=0.4109; Time: F_3.693, 51.70_=5.217, p=0.0017; Interaction: F_3.693, 51.70_ = 1.835, p=0.1409).

Female mice treated prophylactically with vehicle during paclitaxel regimen showed decreased mechanical paw withdrawal thresholds and increased responsiveness to acetone (BL-Day 28). Treatment with CBD attenuated the development of paclitaxel-induced mechanical hypersensitivity in WT female mice, compared to the vehicle-treated group (**Figure 7B**: Treatment: F_1, 14_ = 8.948, p=0.0097; Time: F_4.150, 58.10_=0.4151, p=0.8038; Interaction: F_4.150, 58.10_ = 1.804, p=0.1381). Moreover, CBD attenuated the development of paclitaxel-induced cold hypersensitivity over time in WT female mice (**Figure 7D**: Treatment: F_1, 14_ = 12.58, p=0.0032; Time: F_4.101, 57.42_=5.902, p=0.0004; Interaction: F_4.101, 57.42_ = 1.167, p=0.3351). In paclitaxel treated NAPE-PLD KO female mice, paw withdrawal threshold and cold responsiveness changed across time, irrespective of drug treatment (**Figure 7C**: Treatment: F_1, 14_ = 2.551, p=0.1325; Time: F_4.883, 68.37_=3.898, p=0.0039; Interaction: F_4.883, 68.37_ = 0.9260, p=0.4680; **Figure 7E**: Treatment: F_1, 14_ = 3.308, p=0.0904; Time: F_3.360, 51.23_=9.610, p<0.0001; Interaction: F_3.660, 51.23_ = 0.7008, p=0.5829).

### 3.6 Chronic treatment with CBD attenuates the development of paclitaxel-induced hypersensitivity in both wild-type and GPR55 knock out mice

Paclitaxel-treated male WT (**Figure 8B**) and GPR55 KO mice (**Figure 8C**) treated with vehicle developed mechanical hypersensitivity starting 7 days after the beginning of injections. CBD treatment increased paw withdrawal thresholds, paw withdrawal thresholds changed across time and the interaction between time and treatment was significant in male WT mice (**Figure 8B**: Treatment: F_1, 14_ = 21.73, p=0.0004; Time: F_3.995, 55.93_=3.256, p=0.0180; Interaction: F_3.995, 55.93_ = 2.790, p=0.0349). The same pattern of effects was also observed in GPR55 KO mice (**Figure 8C**: Treatment: F_1, 13_ = 25.79, p=0.0002; Time: F_4.526, 58.84_=2.566, p=0.0411; Interaction: F_4.526, 58.84_ = 2.972, p=0.0219). Prophylactic CBD treatment (10 mg/kg, i.p; 7 daily injections) prevented the development of mechanical hypersensitivity in both WT (p<0.0464 for day 7 to 18 and day 27, otherwise p>0.05, **Figure 8B**) and GPR55 KO mice, when compared to their respective vehicle treatment groups (p<0.0446 for day 11 to 18, otherwise p>0.05, **Figure 8C**).

Prophylactic treatment with CBD also showed a tendency to attenuate paclitaxel-induced cold hypersensitivity in the same cohort of WT mice in a time-dependent manner (**Figure 8D**: Treatment: F_1, 14_ = 4.181, p=0.0601; Time: F_2.430, 34.02_=5.726, p=0.0047; Interaction: F_2.430, 34.02_ = 1.524, p=0.2301). In addition, CBD treatment effectively attenuated the development of cold hypersensitivity in GPR55 KO male mice in a time-dependent manner (**Figure 8E**: Treatment: F_1, 13_ = 6.100, p=0.0281; Time: F_3.953, 51.39_=5.872, p=0.0006; Interaction: F_3.953, 51.39_ = 0.9872, p=0.4222).

## 4. Discussion

Current available treatments for neuropathic pain remain unsatisfactory and cannabinoids have surfaced as potentially promising agents for pain management [24]. CBD, one of the main phytocannabinoids found in *Cannabis* plant, effectively suppresses mechanical and/or thermal hypersensitivity in models of neuropathic pain including that induced by chronic constriction injury of the sciatic nerve, streptozotocin-induced diabetic neuropathy, as well as paclitaxel-induced neuropathic nociception [6,8,9,25–33]. A complex polypharmacology is implicated in the antinociceptive effect of CBD, as suggested by a partial involvement of different molecular targets in the antinociceptive effect of CBD [6–10,13,34]. Here we show that the antinociceptive effects of CBD are dependent upon the enzyme NAPE-PLD in a model of CIPN in mice. In our studies, CBD, administered acutely, attenuated paclitaxel-induced mechanical and cold hypersensitivities, in a dose-dependent manner. CBD administration during the development and maintenance phases of CIPN attenuated hypersensitivities in wild-type, but not NAPE-PLD KO mice. By contrast, the antinociceptive effect of CBD was preserved in GPR55 KO mice in both the development and maintenance phases of paclitaxel-induced hypersensitivity.

N-acylethanolamines (NAEs), the products of NAPE-PLD-mediated N-acylphosphatidylethanolamine (NAPE) hydrolysis, represent a variety of lipid ligands and mediators including G-protein coupled receptors (e.g. endocannabinoid receptors and GPR55), ion channels (e.g. TRPV1), serotonin receptor (5-HT1A) antagonists, ligand activated transcription factors (e.g. peroxisome proliferator receptor-α and peroxisome proliferator receptor-γ) (For review see [35]). Many of these targets are also implicated in the antinociceptive effect of CBD in various pre-clinical models of inflammatory and neuropathic pain [6–11,26,28,34]. In our study, acute CBD administration, in increasing doses given in a within subject regimen, attenuated mechanical and cold hypersensitivity during the maintenance phase of paclitaxel-induced neuropathic pain in mice. To further explore the pharmacological specificity of this acute antinociceptive effect of CBD, different receptor antagonists were administered before CBD treatment in paclitaxel-treated mice. Treatment with CB1 receptor antagonist (AM251) and CB2 receptor antagonist (AM630) did not block the antinociceptive effect of CBD. These observations are consistent with previous reports showing that the antinociceptive effect of CBD are not blocked by cannabinoid receptor antagonists in different pathological pain models, as well as the absence of appreciable binding affinity of CBD to CB1 and CB2 receptors [8,10,36–38].

PPAR receptor isoforms α and γ mediate metabolic and anti-inflammatory effects and endocannabinoid ligands seem to exert significant effects by PPAR receptor activation (for review see [39]). Corroborating in vitro findings, studies in vivo showed that a fluorinated compound derivative of CBD (PECS-101) attenuated mechanical and cold hypersensitivity in chemotherapy-induced neuropathy in mice, an effect mediated by PPAR receptors [13]. In fact, CBD has been shown to act as an agonist at PPARγ receptors, whereas there is comparatively less evidence supporting the activation of PPARα by phytocannabinoids as a class (for review see [40]). Our findings indicate that treatment with PPARα receptor antagonist blocked the antinociceptive effect of CBD over mechanical and cold hypersensitivity, whereas a PPAR γ only attenuated the effect of CBD over mechanical hypersensitivity. Previous findings show that CBD can attenuate neuropathic pain-induced by spinal nerve ligation through a mechanism that required activation of spinal PPAR γ receptors [25,39]. Taken together, these findings suggest that CBD produces acute anti-allodynic effects via a mechanism that likely involves the NAPE-PLD enzyme and PPAR receptors in neuropathic pain models. Moreover, the specific PPAR receptor subtype mediating this antinociceptive effect may differ across pain models. Nonetheless, our study did not specifically evaluate a direct correlation between these two targets (NAPE-PLD, PPAR) amid CBD treatment and its anti-allodynic effect.

Experimental models of CIPN alter endocannabinoid (AEA and 2-AG) levels in multiple tissues, an effect that correlates to development of mechanical and thermal hyperalgesia [41,42]. In addition, the upregulation of NAEs after CBD treatment (3 mg/kg) was found in brain regions (e.g. hippocampus, cerebellum, thalamus, cortex, midbrain, and brainstem) of WT, but NAPE-PLD KO mice [16]. Some of these regions are part of pathways common to CIPN symptom development and go through hyperactivity over time (for review see [43]). In addition, nanoformulated CBD has shown to rapidly suppress neuropathic pain and somatosensory hyperactivity in thalamus, cortex and spinal cord [31]. To further investigate the role of NAPE-PLD in the anti-allodynic effect of CBD, we tested the efficacy of CBD, given during the maintenance phase and during the development phase of paclitaxel-induced mechanical and cold hypersensitivity, in WT and NAPE-PLD KO mice of both sexes. In the maintenance phase, treatment with CBD attenuated mechanical and cold hypersensitivity in WT mice of both sexes, but not in NAPE-PLD KO mice. This is the first report of the involvement of NAPE-PLD in the antinociceptive effect of CBD in a mouse model of neuropathic pain. Previous reports have indicated that CBD treatment can increase expression of NAPE-PLD in brain regions such as the VTA and pre-frontal cortex, in models of neuropsychiatric conditions such as chronic stress and drug addiction, where NAPE-PLD are diminished [44,45]. Additionally, CBD has substantial impact in attenuation of lipid peroxidation, including in neuropathic pain conditions [30,46], as well as modulation of the lipidome in the brain, leading to increase levels of N-acyl ethanolamines with known antinociceptive effects in pre-clinical models(e.g. OEA, AEA)[16,32,47,48], and decrease in inflammatory mediators (e.g.PGE2, PGF2α, TNF)[16,32,46,49].

Treatment with CBD at 10 mg/kg significantly prevented the development of mechanical and cold hypersensitivity in WT male and female mice, but not in NAPE-PLD KO mice of both sexes. Previous studies have demonstrated that CBD attenuates paclitaxel-induced hypersensitivity during its development phase [9,10,32], as well as chronic inflammatory pain [29], but this is the first report to show that this antinociceptive effect is abrogated in NAPE-PLD KO mice. Although we observed efficacy of CBD during the development phase of paclitaxel-induced hypersensitivity in both sexes, the efficacy observed in female mice appears to be mild, whereas the effect in males is robust. There is limited literature on sex differences in response to CBD, but few studies have reported mixed effects of females to CBD treatment in experimental models evaluating affective responses, such as anxiety, depression, conditioned avoidance, and aversion, as well as memory and locomotor activity [50–52]. In fact, efficacy of CBD in panic-like response of mice has been shown to be dependent of a specific estrous cycle phase in female mice [53]. Pharmacokinetic results indicate that CBD treatment in females leads to higher CBD concentration in specific tissues, when compared to males [54]. In addition, while there is a higher early plasma concentration of CBD and early formation of CBD metabolites in female mice, the volume of distribution is more than 2-fold higher in male mice [55]. Combined treatment of CBD and paclitaxel could increase the efficacy of paclitaxel, leading to simultaneous increase in adverse events [56]. Nonetheless, the chemotherapy efficacy of paclitaxel has been shown unaltered by CBD treatment in the viability of breast cancer cells [10]. Further studies are required to investigate the sex differences found in the preventive effect of CBD, as well as a potential role of estrous cycle in the efficacy of CBD during the development phase of paclitaxel-induced hypersensitivity in female mice.

CBD has been successfully used in clinical trials as an anti-seizure treatment in severe and refractory seizure syndromes such as Dravet syndrome and Lennox-Gastaut syndrome [57]. Recent studies have implicated GPR55, a de-orphanized g-protein couple receptor, as a target for the seizure-reducing properties of CBD. CBD acts as GPR55 antagonist and dose-dependently reduced pentylenetetrazole-induced seizures in mice, an effect that was absent in GPR55 KO mice [22]. Since GPR55 has been associated with the establishment of hyperalgesia in models of inflammatory and neuropathic pain [58] and CBD has been validated as an antagonist of GPR55 in vivo [22], the current study also tested whether GPR55 KO mice would respond differently to CBD treatment during the development and maintenance phases of paclitaxel-induced mechanical and cold hypersensitivity. CBD was efficacious in preventing and reversing paclitaxel-induced mechanical and cold hypersensitivity in both WT and GPR55-KO mice. Taken together, these findings suggest that any association between CBD’s anti-allodynic effects and GPR55 receptors in paclitaxel-induced neuropathic pain is minimal or absent, while reinforcing the role of NAPE-PLD in mediating CBD’s anti-allodynic actions across diverse models of pathological pain.

In conclusion, our results further elucidate the pharmacological mechanism involved in the antinociceptive effect of CBD, indicating that NAPE-PLD plays a pivotal role in both prevention and reversal of hypersensitivity by CBD treatment in chemotherapy-induced neuropathic pain. Considering that available pharmacotherapies used for the management of chemotherapy-induced neuropathic pain fail to produce satisfactory symptom relief, CBD could emerge as an effective treatment for these conditions. More work is necessary to address sex differences found in response to CBD and to evaluate the pathways that potentially engage NAPE-PLD enzyme upon CBD treatment in experimental neuropathic pain conditions.

## Acknowledgements

The authors are grateful to Heather Bradshaw, Craig Farris, Daniele Piomelli and Mario van der Stelt for helpful discussions during the development of the study.

The authors confirm that the data supporting the findings of this study are uploaded in Zenodo data repository: https://doi.org/10.5281/zenodo.20562653.

## References

[1] D. Bouhassira, Neuropathic pain: Definition, assessment and epidemiology, Rev. Neurol. (Paris). 175 (2019) 16–25. 10.1016/J.NEUROL.2018.09.016.

[2] N.B. Finnerup, R. Kuner, T.S. Jensen, Neuropathic Pain: From Mechanisms to Treatment, Physiol. Rev. 101 (2021) 259–301. 10.1152/PHYSREV.00045.2019.

[3] G.K. Silva-Cardoso, C.R.A. Leite-Panissi, Chronic Pain and Cannabidiol in Animal Models: Behavioral Pharmacology and Future Perspectives, Cannabis Cannabinoid Res. 8 (2023). 10.1089/CAN.2022.0096.

[4] J. Peng, M. Fan, C. An, F. Ni, W. Huang, J. Luo, A narrative review of molecular mechanism and therapeutic effect of cannabidiol (CBD), Basic Clin. Pharmacol. Toxicol. 130 (2022) 439–456. 10.1111/BCPT.13710.

[5] C.H.A. Jesus, M.V. Ferreira, A.T. Gasparin, E.S. Rosa, K. Genaro, J.A. de S. Crippa, J.G. Chichorro, J.M. da Cunha, Cannabidiol enhances the antinociceptive effects of morphine and attenuates opioid-induced tolerance in the chronic constriction injury model, Behavioural Brain Research 435 (2022). 10.1016/J.BBR.2022.114076.

[6] G.K. Silva-Cardoso, W. Lazarini-Lopes, J.E. Hallak, J.A. Crippa, A.W. Zuardi, N. Garcia-Cairasco, C.R.A. Leite-Panissi, Cannabidiol effectively reverses mechanical and thermal allodynia, hyperalgesia, and anxious behaviors in a neuropathic pain model: Possible role of CB1 and TRPV1 receptors, Neuropharmacology 197 (2021). 10.1016/J.NEUROPHARM.2021.108712.

[7] C.C. Toth, N.M. Jedrzejewski, C.L. Ellis, W.H. Frey, Cannabinoid-mediated modulation of neuropathic pain and microglial accumulation in a model of murine type I diabetic peripheral neuropathic pain, Mol. Pain 6 (2010). 10.1186/1744-8069-6-16.

[8] C.H.A. Jesus, D.D.B. Redivo, A.T. Gasparin, B.B. Sotomaior, M.C. de Carvalho, K. Genaro, A.W. Zuardi, J.E.C. Hallak, J.A. Crippa, J.M. Zanoveli, J.M. da Cunha, Cannabidiol attenuates mechanical allodynia in streptozotocin-induced diabetic rats via serotonergic system activation through 5-HT1A receptors, Brain Res. 1715 (2019) 156–164. 10.1016/J.BRAINRES.2019.03.014.

[9] S.J. Ward, M.D. Ramirez, H. Neelakantan, E.A. Walker, Cannabidiol Prevents the Development of Cold and Mechanical Allodynia in Paclitaxel-Treated Female C57Bl6 Mice, Anesth. Analg. 113 (2011) 947. 10.1213/ANE.0B013E3182283486.

[10] S.J. Ward, S.D. McAllister, R. Kawamura, R. Murase, H. Neelakantan, E.A. Walker, Cannabidiol inhibits paclitaxel-induced neuropathic pain through 5-HT(1A) receptors without diminishing nervous system function or chemotherapy efficacy, Br. J. Pharmacol. 171 (2014) 636–645. 10.1111/BPH.12439.

[11] J. Mlost, M. Bryk, K. Starowicz, Cannabidiol for Pain Treatment: Focus on Pharmacology and Mechanism of Action, Int. J. Mol. Sci. 21 (2020) 1–22. 10.3390/IJMS21228870.

[12] M. Tham, O. Yilmaz, M. Alaverdashvili, M.E.M. Kelly, E.M. Denovan-Wright, R.B. Laprairie, Allosteric and orthosteric pharmacology of cannabidiol and cannabidiol-dimethylheptyl at the type 1 and type 2 cannabinoid receptors, Br. J. Pharmacol. 176 (2019) 1455. 10.1111/BPH.14440.

[13] N.R. Silva, F.I.F. Gomes, A.H.P. Lopes, I.L. Cortez, J.C. dos Santos, C.E.A. Silva, R. Mechoulam, F.V. Gomes, T.M. Cunha, F.S. Guimarães, The Cannabidiol Analog PECS-101 Prevents Chemotherapy-Induced Neuropathic Pain via PPARγ Receptors, Neurotherapeutics 19 (2022) 434–449. 10.1007/S13311-021-01164-W/FIGURES/6.

[14] R.A. Gray, B.J. Whalley, The proposed mechanisms of action of CBD in epilepsy, Epileptic Disorders 22 (2020) S10–S15. 10.1684/EPD.2020.1135.

[15] S. Armin, S. Muenster, M. Abood, K. Benamar, GPR55 in the brain and chronic neuropathic pain, Behavioural Brain Research 406 (2021). 10.1016/J.BBR.2021.113248.

[16] E. Leishman, M. Manchanda, R. Thelen, S. Miller, K. Mackie, H.B. Bradshaw, Cannabidiol’s Upregulation of N-acyl Ethanolamines in the Central Nervous System Requires N-acyl Phosphatidyl Ethanolamine-Specific Phospholipase D, Cannabis Cannabinoid Res. 3 (2018) 228–241. 10.1089/CAN.2018.0031/SUPPL_FILE/SUPP_DATA.DOCX.

[17] M. Zimmermann, Ethical guidelines for investigations of experimental pain in conscious animals, Pain 16 (1983) 109–110. 10.1016/0304-3959(83)90201-4.

[18] X. Lin, A.S. Dhopeshwarkar, M. Huibregtse, K. MacKie, A.G. Hohmann, Slowly signaling G protein-biased CB2 cannabinoid receptor agonist LY2828360 suppresses neuropathic pain with sustained efficacy and attenuates morphine tolerance and dependence, Mol. Pharmacol. 93 (2018) 49–62. 10.1124/MOL.117.109355/-/DC1.

[19] L. Deng, J. Guindon, B.L. Cornett, A. Makriyannis, K. Mackie, A.G. Hohmann, Chronic cannabinoid CB2 activation reverses paclitaxel neuropathy without tolerance or CB1-dependent withdrawal, Biol. Psychiatry 77 (2014) 475. 10.1016/J.BIOPSYCH.2014.04.009.

[20] K.G. Guenther, X. Lin, Z. Xu, A. Makriyannis, J. Romero, C.J. Hillard, K. Mackie, A.G. Hohmann, Cannabinoid CB2 receptors in primary sensory neurons are implicated in CB2 agonist-mediated suppression of paclitaxel-induced neuropathic nociception and sexually-dimorphic sparing of morphine tolerance, Biomedicine and Pharmacotherapy 176 (2024). 10.1016/j.biopha.2024.116879.

[21] J.L. Wirt, L.A. Ferreira, C.H. Alves Jesus, T.J. Woodward, I. Oliva, Z. Xu, J.D. Crystal, R.H. Pepin, R.B. Silverman, A.G. Hohmann, Efficacy of GABA aminotransferase inactivator OV329 in models of neuropathic and inflammatory pain without tolerance or addiction, Proc. Natl. Acad. Sci. U. S. A. 122 (2025). 10.1073/PNAS.2318833121.

[22] E.C. Rosenberg, S. Chamberland, M. Bazelot, E.R. Nebet, X. Wang, S. McKenzie, S. Jain, S. Greenhill, M. Wilson, N. Marley, A. Salah, S. Bailey, P.H. Patra, R. Rose, N. Chenouard, S.D. Sun, D. Jones, G. Buzsáki, O. Devinsky, G. Woodhall, H.E. Scharfman, B.J. Whalley, R.W. Tsien, Cannabidiol modulates excitatory-inhibitory ratio to counter hippocampal hyperactivity, Neuron 111 (2023) 1282–1300.e8. 10.1016/j.neuron.2023.01.018.

[23] O. Sasso, G. Moreno-Sanz, C. Martucci, N. Realini, M. Dionisi, L. Mengatto, A. Duranti, G. Tarozzo, G. Tarzia, M. Mor, R. Bertorelli, A. Reggiani, D. Piomelli, Antinociceptive effects of the N-acylethanolamine acid amidase inhibitor ARN077 in rodent pain models, Pain 154 (2013) 350–360. 10.1016/J.PAIN.2012.10.018.

[24] B.W. Johnson, N.H. Strand, J.C. Raynak, C. Jara, K. Habtegiorgis, B.A. Hand, S. Hong, J.A. Maloney, Cannabinoids in Chronic Pain Management: A Review of the History, Efficacy, Applications, and Risks, Biomedicines 13 (2025). 10.3390/BIOMEDICINES13030530.

[25] A.M. Islas-Espinoza, I.I. Ramos-Rodríguez, M.J. Escoto-Rosales, J.M. Pizaña-Encarnación, D.K. Morales-Galindo, N.L. Caram-Salas, M. Déciga-Campos, E.J. Rodríguez-Palma, V. Granados-Soto, Cannabidiol reduces neuropathic pain and cognitive impairments through activation of spinal PPARγ, Journal of Pain 30 (2025) 105378. 10.1016/j.jpain.2025.105378.

[26] D. De Gregorio, R.J. McLaughlin, L. Posa, R. Ochoa-Sanchez, J. Enns, M. Lopez-Canul, M. Aboud, S. Maione, S. Comai, G. Gobbi, Cannabidiol modulates serotonergic transmission and reverses both allodynia and anxiety-like behavior in a model of neuropathic pain, Pain 160 (2019) 136–150. 10.1097/J.PAIN.0000000000001386.

[27] D.L. de Almeida, R.C. Mendes Ferreira, F.C. Fonseca, D.P. Dias Machado, D.D. Aguiar, F.S. Guimaraes, I.D.G. Duarte, T.R.L. Romero, Cannabidiol induces systemic analgesia through activation of the PI3Kγ/nNOS/NO/KATP signaling pathway in neuropathic mice. A KATP channel S-nitrosylation-dependent mechanism, Nitric Oxide 146 (2024) 1–9. 10.1016/J.NIOX.2024.02.005.

[28] K.M. King, A.M. Myers, A.J. Soroka-Monzo, R.F. Tuma, R.J. Tallarida, E.A. Walker, S.J. Ward, Single and combined effects of Δ9-tetrahydrocannabinol and cannabidiol in a mouse model of chemotherapy-induced neuropathic pain, Br. J. Pharmacol. 174 (2017) 2832–2841. 10.1111/BPH.13887;WEBSITE:WEBSITE:BPSPUBS;WGROUP:STRING:PUBLICATION.

[29] H.T. Philpott, M. O’Brien, J.J. McDougall, Attenuation of early phase inflammation by cannabidiol prevents pain and nerve damage in rat osteoarthritis, Pain 158 (2017) 2442–2451. 10.1097/J.PAIN.0000000000001052.

[30] V. Baron-Flores, A. Diaz-Ruiz, J. Manzanares, C. Rios, M. Burelo, G. Jardon-Guadarrama, M. de los Á. Martínez-Cárdenas, A. Mata-Bermudez, Cannabidiol attenuates hypersensitivity and oxidative stress after traumatic spinal cord injury in rats, Neurosci. Lett. 788 (2022). 10.1016/j.neulet.2022.136855.

[31] J. Feng, J. Page, L. Chung, Z. He, K.H. Wang, Rapid suppression of neuropathic pain and somatosensory hyperactivity by nano-formulated cannabidiol, Cell Chem. Biol. 32 (2025) 1412–1428.e5. 10.1016/J.CHEMBIOL.2025.10.005.

[32] R. dos Santos, F. Veras, G. Netto, L. Elisei, C. Sorgi, L. Faccioli, G. Galdino, Cannabidiol prevents chemotherapy-induced neuropathic pain by modulating spinal TLR4 via endocannabinoid system activation, Journal of Pharmacy and Pharmacology 75 (2023) 655–665. 10.1093/JPP/RGAD023.

[33] D.D. Aguiar, C. da Costa Oliveira, F.C.S. Fonseca, D.L. de Almeida, W.V. Campos Pereira, F.S. Guimarães, A.C. Perez, I.D.G. Duarte, T.R.L. Romero, Peripherally injected canabidiol reduces neuropathic pain in mice: Role of the 5-HT1A and TRPV1 receptors, Biochem. Biophys. Res. Commun. 660 (2023) 58–64. 10.1016/J.BBRC.2023.04.022.

[34] B. Costa, G. Giagnoni, C. Franke, A.E. Trovato, M. Colleoni, Vanilloid TRPV1 receptor mediates the antihyperalgesic effect of the nonpsychoactive cannabinoid, cannabidiol, in a rat model of acute inflammation, Br. J. Pharmacol. 143 (2004) 247. 10.1038/SJ.BJP.0705920.

[35] E.D. Mock, B. Gagestein, M. van der Stelt, Anandamide and other N-acylethanolamines: A class of signaling lipids with therapeutic opportunities, Prog. Lipid Res. 89 (2023). 10.1016/j.plipres.2022.101194.

[36] T. Bisogno, L. HanusAE, L. De Petrocellis, S. Tchilibon, D.E. Ponde, I. Brandi, A. Schiano Moriello, J.B. Davis, R. Mechoulam, V. Di Marzo, Molecular targets for cannabidiol and its synthetic analogues: effect on vanilloid VR1 receptors and on the cellular uptake and enzymatic hydrolysis of anandamide, Br. J. Pharmacol. 134 (2001) 852. www.nature.com/bjp (accessed June 8, 2022).

[37] J. Bouma, J.D. Broekhuis, C. van der Horst, P. Kumar, A. Ligresti, M. van der Stelt, L.H. Heitman, Dual allosteric and orthosteric pharmacology of synthetic analog cannabidiol-dimethylheptyl, but not cannabidiol, on the cannabinoid CB2 receptor, Biochem. Pharmacol. 218 (2023). 10.1016/j.bcp.2023.115924.

[38] N. Schuelert, J.J. McDougall, The abnormal cannabidiol analogue O-1602 reduces nociception in a rat model of acute arthritis via the putative cannabinoid receptor GPR55, Neurosci. Lett. 500 (2011) 72–76. 10.1016/J.NEULET.2011.06.004.

[39] F.A. Iannotti, R.M. Vitale, The Endocannabinoid System and PPARs: Focus on Their Signalling Crosstalk, Action and Transcriptional Regulation, Cells 10 (2021) 586. 10.3390/CELLS10030586.

[40] S.E. O’Sullivan, An update on PPAR activation by cannabinoids, Br. J. Pharmacol. 173 (2016) 1899. 10.1111/BPH.13497.

[41] I.A. Khasabova, S. Khasabov, J. Paz, C. Harding-Rose, D.A. Simone, V.S. Seybold, Cannabinoid Type-1 Receptor Reduces Pain and Neurotoxicity Produced by Chemotherapy, The Journal of Neuroscience 32 (2012) 7091. 10.1523/JNEUROSCI.0403-12.2012.

[42] I.A. Khasabova, X. Yao, J. Paz, C.T. Lewandowski, A.E. Lindberg, L. Coicou, N. Burlakova, D.A. Simone, V.S. Seybold, JZL184 is anti-hyperalgesic in a murine model of cisplatin-induced peripheral neuropathy, Pharmacol. Res. 90 (2014) 67–75. 10.1016/J.PHRS.2014.09.008.

[43] M. Omran, E.K. Belcher, N.A. Mohile, S.R. Kesler, M.C. Janelsins, A.G. Hohmann, I.R. Kleckner, Review of the Role of the Brain in Chemotherapy-Induced Peripheral Neuropathy, Front. Mol. Biosci. 8 (2021) 693133. 10.3389/FMOLB.2021.693133.

[44] F.F. Scarante, V.D. Lopes, E.J. Fusse, M.A. Vicente, G.H.D. de Abreu, V. Nardini, C.A. Sorgi, P.H.C. Lirio, F. Guo, H.C. Lu, J.E.C. Hallak, J.A. Crippa, L.H. Faccioli, K.U. Sales, F.S. Guimarães, K. Mackie, A.C. Campos, Cannabidiol reduces the latency for the behavioral effect of escitalopram in chronically stressed male mice: involvement of NAPE-PLD expressed in parvalbumin-positive interneurons and the prefrontal cortex, Neuropharmacology 291 (2026). 10.1016/J.NEUROPHARM.2026.110894.

[45] V.G. Metz, J.L.O. da Rosa, D.R. Rossato, M.E. Burger, C.S. Pase, Cannabidiol treatment prevents drug reinstatement and the molecular alterations evoked by amphetamine on receptors and enzymes from dopaminergic and endocannabinoid systems in rats, Pharmacol. Biochem. Behav. 218 (2022). 10.1016/j.pbb.2022.173427.

[46] B. Costa, A.E. Trovato, F. Comelli, G. Giagnoni, M. Colleoni, The non-psychoactive cannabis constituent cannabidiol is an orally effective therapeutic agent in rat chronic inflammatory and neuropathic pain, Eur. J. Pharmacol. 556 (2007) 75–83. 10.1016/J.EJPHAR.2006.11.006.

[47] M. Suardíaz, G. Estivill-Torrús, C. Goicoechea, A. Bilbao, F. Rodríguez de Fonseca, Analgesic properties of oleoylethanolamide (OEA) in visceral and inflammatory pain, Pain 133 (2007) 99–110. 10.1016/J.PAIN.2007.03.008.

[48] A.K. Schreiber, M. Neufeld, C.H.A. Jesus, J.M. Cunha, Peripheral antinociceptive effect of anandamide and drugs that affect the endocannabinoid system on the formalin test in normal and streptozotocin-diabetic rats, Neuropharmacology 63 (2012) 1286–1297. 10.1016/J.NEUROPHARM.2012.08.009.

[49] T. Lowin, R. Tingting, J. Zurmahr, T. Classen, M. Schneider, G. Pongratz, Cannabidiol (CBD): a killer for inflammatory rheumatoid arthritis synovial fibroblasts, Cell Death & Disease 2020 11:8 11 (2020) 714-. 10.1038/s41419-020-02892-1.

[50] J.S. Kaplan, J.K. Wagner, K. Reid, F. McGuinness, S. Arvila, M. Brooks, H. Stevenson, J. Jones, B. Risch, T. McGillis, R. Budinich, E. Gambell, B. Predovich, Cannabidiol Exposure During the Mouse Adolescent Period Is Without Harmful Behavioral Effects on Locomotor Activity, Anxiety, and Spatial Memory, Front. Behav. Neurosci. 15 (2021). 10.3389/FNBEH.2021.711639.

[51] S. Ledesma-Corvi, E. Hernández-Hernández, M.J. García-Fuster, Exploring pharmacological options for adolescent depression: a preclinical evaluation with a sex perspective, Transl. Psychiatry 12 (2022). 10.1038/S41398-022-01994-Y.

[52] I.D.S. Maciel, G.H.D.D. Abreu, C.T. Johnson, R. Bonday, H.B. Bradshaw, K. Mackie, H.C. Lu, Perinatal CBD or THC Exposure Results in Lasting Resistance to Fluoxetine in the Forced Swim Test: Reversal by Fatty Acid Amide Hydrolase Inhibition, Cannabis Cannabinoid Res. 7 (2022) 318–327. 10.1089/CAN.2021.0015.

[53] P.M. Hernandes, M.F. Batistela, J.M. Nascimento-Silva, A.T. Frias, M. Matthiesen, A.C. Campos, T.A. Lovick, H. Zangrossi, Sex and estrous cycle-linked differences in the effect of cannabidiol on panic-like responding in rats and mice, Behavioural Brain Research 455 (2023) 114663. 10.1016/J.BBR.2023.114663.

[54] R.B. Child, M.J. Tallon, Cannabidiol (CBD) Dosing: Plasma Pharmacokinetics and Effects on Accumulation in Skeletal Muscle, Liver and Adipose Tissue, Nutrients 14 (2022). 10.3390/NU14102101.

[55] M.E. Olawale, M.D. Lad, M.C. Anyachebelu, S. Luo, P. Lazarus, Sex differences in the disposition of cannabidiol and its metabolites in mice, Journal of Cannabis Research 2026 (2026). 10.1186/S42238-026-00427-7.

[56] J.D. Brown, A.G. Winterstein, Potential Adverse Drug Events and Drug–Drug Interactions with Medical and Consumer Cannabidiol (CBD) Use, J. Clin. Med. 8 (2019). 10.3390/JCM8070989.

[57] A.T. Berg, T. Dixon-Salazar, M.A. Meskis, S.R. Danese, N.M.D. Le, M.S. Perry, Caregiver-reported outcomes with real-world use of cannabidiol in Lennox-Gastaut syndrome and Dravet syndrome from the BECOME survey, Epilepsy Res. 200 (2023) 107280. 10.1016/J.EPLEPSYRES.2023.107280.

[58] P.C. Staton, J.P. Hatcher, D.J. Walker, A.D. Morrison, E.M. Shapland, J.P. Hughes, E. Chong, P.K. Mander, P.J. Green, A. Billinton, M. Fulleylove, H.C. Lancaster, J.C. Smith, L.T. Bailey, A. Wise, A.J. Brown, J.C. Richardson, I.P. Chessell, The putative cannabinoid receptor GPR55 plays a role in mechanical hyperalgesia associated with inflammatory and neuropathic pain, Pain 139 (2008) 225–236. 10.1016/J.PAIN.2008.04.006.

